# A Nerve-Fibroblast Axis in Mammalian Lung Fibrosis

**DOI:** 10.1101/2024.09.09.611003

**Authors:** Genta Ishikawa, Xueyan Peng, Alexander Ghincea, John McGovern, Jana Zielonka, Advait Jeevanandam, Shuai Shao, Samuel Woo, Daisuke Okuno, Sheeline Yu, Chris J. Lee, Angela Liu, Tina Saber, Buqu Hu, Ying Sun, Ruijuan Gao, Karam Al Jumaily, Robert Homer, Monique Hinchcliff, Carol Feghali-Bostwick, Tomokazu S. Sumida, Maor Sauler, Jose L. Gomez, Huanxing Sun, Changwan Ryu, Erica L. Herzog

**Affiliations:** Department of Internal Medicine, Section of Pulmonary, Critical Care, and Sleep Medicine, School of Medicine, Yale University, New Haven, CT, USA; Department of Immunobiology, School of Medicine, Yale University, New Haven, CT, USA; Institute of Medicinal Biotechnology, Chinese Academy of Medical Sciences and Peking Union Medical College, No.1 Tiantan Xili, Beijing 100050, China; Department of Pathology, School of Medicine, Yale University, New Haven, CT, USA; Department of Internal Medicine, Section of Rheumatology, Allergy and Immunology, School of Medicine, Yale University, New Haven, CT, USA; Department of Medicine, Division of Rheumatology and Immunology, Medical University of South Carolina, SC, USA; Department of Neurology, School of Medicine, Yale University, New Haven, CT, USA

## Abstract

Fibrosis contributes to incurable pathologies in vital organs including the lung. Myofibroblasts are fibrogenic effector cells that accumulate via incompletely understood mechanisms. We discovered that α1-adrenoreceptor expressing myofibroblasts receive sympathetic nerve-derived noradrenergic inputs in fibrotic mouse and human lungs. We combined optical clearing, whole lung imaging, cell-specific gene deletion in sympathetic nerves and myofibroblasts, pharmacologic interventions, sympathetic nerve co-culture and precision-cut lung slices, with analysis of bronchoalveolar lavage fluid, lung tissues, single-cell RNA sequencing datasets, and isolated lung fibroblasts from patients with diverse forms of pulmonary fibrosis to characterize a fibrogenic unit comprised of aberrantly patterned sympathetic nerves and α1-adrenoreceptor subtype D expressing myofibroblasts. The discovery of this previously undefined nerve-fibroblast axis that is conserved across species demonstrates the pivotal contribution of nerves to tissue remodeling and heralds a novel paradigm in fibrosis research.

## INTRODUCTION

Tissue fibrosis arising from ineffective wound healing is implicated in human pathologies affecting critical organs such as the lung^1^, heart^2^, liver^3^, kidney^4^, and skin^5^. The process is currently viewed as irreversible and contributes to upwards of 45% deaths in the industrialized world^6^. While in the last two decades progress has been made with the development of modestly effective therapies for pulmonary fibrosis^7,8^, curative treatments are lacking and the disease burden remains high. This shortcoming results in part from limited understanding of cellular and molecular mechanisms contributing to pathology and presents an imperative for the discovery of new opportunities for research and therapeutic innovation.

Successful wound repair requires the expansion of activated myofibroblasts that receive and respond to microenvironmental signals by contracting the wound bed and producing extracellular matrix^9^. From a tissue perspective, repair resolution and functional restoration require that myofibroblast programming shift from activated expansion to quiescent regression^10^. Because myofibroblast persistence is a hallmark of fibrotic diseases affecting the lung and other organs^11^, fibrosis has evolved to be viewed as an unrelenting form of maladaptive repair and regenerative failure. Interestingly, while the mechanisms driving myofibroblast activation have been well studied^10^, less is known about myofibroblast persistence. This pathology is proposed to involve emergent effector cell populations that experience perturbations in critical decisions related to their functional state and ultimate fate. The discovery of fundamental yet intervenable processes governing this biology would be a major advance for the treatment of fibrosis affecting the lung and other organs.

The nervous system controls organismal homeostasis in health and disease and is increasingly implicated in tissue-level injury and remodeling responses^12–18^. In the lung, autonomic input controls the physiology of airways, blood vessels, and secretory glands but there is only limited information regarding the role of local innervation in disease processes affecting alveolar regions.

Several groups, including our own, have studied autonomic and sensory nerves in conditions of inflammatory lung remodeling but most of these studies have focused on immune cell activation, airway disorders, and/or host defense^14,19–21^. Far less is known about the contribution of lung innervation to effector cells such as myofibroblasts under conditions of repair and pathologic remodeling affecting alveoli, which are critical gas exchange regions in the lung. While alveolar myofibroblasts have been recently shown to require autonomic nerve input for proper lung development^22^, an analogous relationship between nerves and the myofibroblasts that arise during alveolar fibrosis is highly plausible but has not been shown. The discovery that the lung’s autonomic nerve supply directs myofibroblast function during alveolar remodeling would substantially advance the understanding of fibrosis pathobiology and provide opportunities for therapeutic intervention.

Using an experimental platform comprised of optical clearing, whole lung imaging, novel mouse lines, sympathetic nerve-specific gene deletion, pharmacologic interventions, sympathetic nerve co-culture, precision-cut lung slices, and analysis of bronchoalveolar lavage fluid, lung cells and tissues, and single-cell RNA sequencing datasets of patients with diverse forms of pulmonary fibrosis, we report a new role for autonomic innervation, specifically sympathetic nerves, in directing myofibroblast biology during alveolar fibrosis. This work provides new areas for therapeutic development and illuminates a previously unrecognized process by which two discrete organ systems – the autonomic nervous system and the lungs – form a functional axis during repair and fibrosis in higher organisms.

## RESULTS

### Aberrant sympathetic nerve patterning in fibrotic lungs

To study whether lung sympathetic innervation directs pulmonary fibrosis, we employed the widely used bleomycin model^23^. This approach consistently induces alveolar fibrosis, evidenced by trichrome staining of lung tissues (Fig. 1A and B) and biochemical collagen measurements (Fig. 1C). This fibrotic response is paralleled by accumulation of local but not circulating noradrenaline (Fig. 1D and E) that may originate from the lung’s sympathetic nerve supply. To more firmly illustrate this concept, we performed optical clearing followed by whole lung tissue imaging using Adipo-Clear protocols^24^ and light-sheet microscopy with three-dimensional reconstruction immunostaining for tyrosine hydroxylase (TH) and α-SMA (Fig. 1F and G). TH is the rate-limiting enzyme in catecholamine synthesis and is used to identify sympathetic nerves in adult organs^25^. Uninjured mouse lungs demonstrated the previously described presence of TH+ nerves in α-SMA+ regions of proximal airways and blood vessels^26^, with a minimal presence in the distal airways (Fig. 1H, 1J-L, S1A and B). Notably, in fibrotic lungs, TH+ nerves displayed aberrant airway patterning, with enhanced signal in proximal regions and physical extension into distal regions (Fig. 1I, 1M-O, S1C and D). In the latter regions they existed in proximity to fibrotic alveolar regions that were enriched for the α-SMA+ parenchymal myofibroblasts that are viewed as pivotal effector cells in fibrotic lung pathologies^27^.

**Fig. 1:**
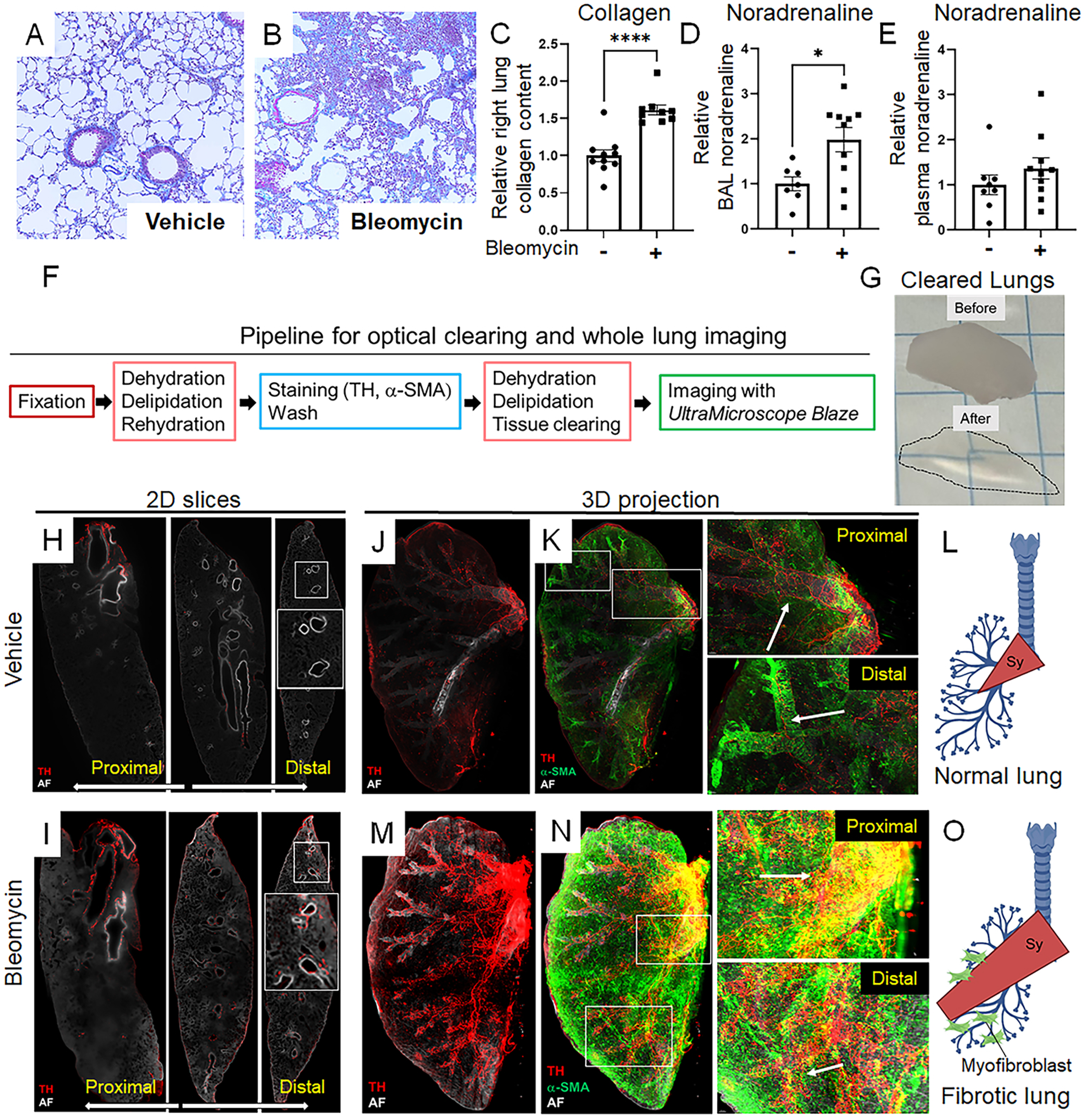
Aberrant sympathetic nerve patterning in fibrotic lungs. (A-E) Wild-type mice received 2.0 U/kg orotracheal vehicle or bleomycin on Day 0 and were sacrificed on Day 14. Bleomycin administration induced alveolar fibrosis, evidenced by trichrome staining (A, B), elevated collagen (C; *P* < 0.0001), and increased noradrenaline concentrations in the BAL (D; *P* = 0.0138), but not plasma (E). (F, G) Experimental pipeline for sympathetic nerve imaging (F) in whole lungs pre/post Adipo-Clear (G). (H-O) Light-sheet microscopy visualized sympathetic innervations and myofibroblasts immunolabeled with anti-TH (red) and anti-α-SMA (green) respectively. 2D projection images with TH signals surrounding the proximal but not distal airways of uninjured lungs (H), while fibrotic lungs showed a notable increase in overall TH signal region extending into peripheral airways (I). (J-L) 3D projections in uninjured lungs demonstrated that TH+ nerves were localized near proximal airways, with minimal extension into distal airways (white arrows). (M-O) In fibrotic lungs, TH+ nerves extended into peripheral airways adjacent to α-SMA+ myofibroblasts (white arrows). Data are presented as mean ± SEM with statistical analyses using Student’s t-test. \**P* < 0.05, \*\*\*\**P* < 0.0001. AF, autofluorescence; α-SMA, alpha-smooth muscle actin; BAL, bronchoalveolar lavage; Sy, sympathetic nerve; TH, tyrosine hydroxylase.

### Sympathetic nerve co-culture amplifies TGFβ1-induced ACTA2 expression in fibroblasts

The identification of TH+ nerves in a tissue milieu characterized by noradrenaline and α-SMA+ myofibroblasts suggests a functional innervation unit in which sympathetic nerves transmit fibrogenic signals to α-SMA+ myofibroblasts in alveolar regions of the adult lung. We tested this hypothesis in a reductive co-culture model of sympathetic nerves and fibroblasts. A well accepted sympathetic nerve culture model^28^ employs primary neuronal cells harvested from neonatal mouse sympathetic chain ganglia (Fig. 2A) in nerve growth factor-containing culture media to allow the development of axonal extensions, TH expression, and noradrenaline production (Fig. 2B-F). To ask whether these cells might impact myofibroblast accumulation, sympathetic nerves were co-cultured with early passage normal human lung fibroblasts (NHLFs) that had been previously isolated via explant outgrowth from parenchyma of donor lungs that were not used for transplantation. Studies were performed in the presence or absence of the profibrotic mediator TGFβ1 (Fig. 2G-J). In this setup, sympathetic nerves alone were insufficient to alter *ACTA2* expression in NHLFs (Fig. 2I). In contrast, under the TGFβ1-rich conditions that exist in fibrotic tissues, NHLFs co-cultured with sympathetic nerves developed substantial increase in the relative expression of *ACTA2* (Fig. 2J). These findings indicate that sympathetic nerves amplify myofibroblast effector properties such as *ACTA2* expression in human lung fibroblasts exposed to fibrotic stimuli.

**Fig. 2:**
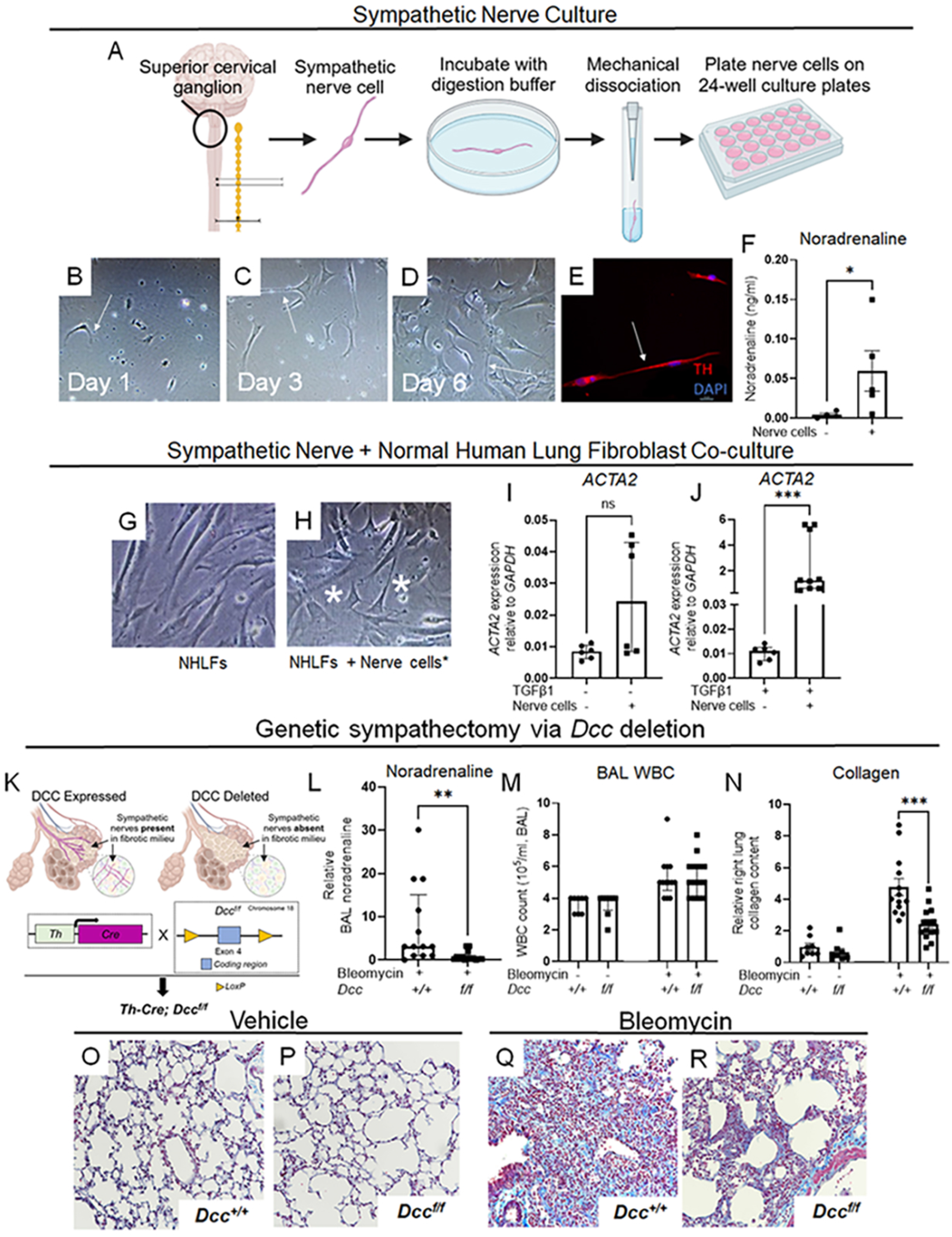
Fibrogenic function of sympathetic neurons *in vitro* and *in vivo*. (A-F) Sympathetic neurons harvested from the superior cervical ganglion of mice at postnatal days 1-3 were cultured in 24-well plates (A), demonstrating time-dependent axonal extension (white arrows, B-D), tyrosine hydroxylase (TH) immunostaining (red, E), and increased noradrenaline concentrations in the culture supernatant by day 3 (F; *P* = 0.0483). (G-J) After 7 days of growth, sympathetic neurons were co-cultured with normal human lung fibroblasts (NHLFs) in the presence or absence of TGFβ1 (5 ng/ml) for 48 hours (nerve cells: white asterisks, G, H). In the absence of TGFβ1, co-culture with nerve cells did not alter *ACTA2* expression in NHLFs (I). In TGFβ1-enriched conditions, co-culture significantly increased *ACTA2* expression in NHLFs (J; *P* = 0.0004). (K-R) Mice with a sympathetic nerves intact (*Th-Cre; Dcc^+/+^)* or deleted (*Th-Cre; Dcc^f/f^*, K) received orotracheal bleomycin or PBS and were sacrificed on Day 14. Following bleomycin, *Th-Cre; Dcc^f/f^* mice displayed reduced BAL noradrenaline (L; *P* = 0.0024), unchanged BAL white blood cell counts (M), reduced collagen accumulation (N; *P* = 0.0002) and improved trichrome staining of alveolar regions (O-R). Images were captured at 20x magnification. Data are presented as mean ± SEM or median ± IQR, with statistical analyses performed using Student’s t-test for normally distributed data and the Mann-Whitney test for non-normally distributed data. \**P* < 0.05, \*\**P* < 0.01, \*\*\**P* < 0.001. DAPI, 4’,6-diamidino-2-phenylindole; DCC, Deleted in Colorectal Cancer; NHLF, normal human lung fibroblast; TH, tyrosine hydroxylase.

### Bleomycin-induced fibrotic endpoints are mitigated following genetic denervation of sympathetic neurons

Next, we asked whether sympathetic nerves are directly required for fibrosis. This inquiry aligns with the hypothesis that effector cells within fibrotic areas receive and respond to neurotransmitters released by nerves in their vicinity. Although there is some evidence suggesting a connection between sympathetic innervation and fibrosis in the lungs and other organs^12–18^, a direct and definitive connection has yet to be established. Here we employed a well-characterized genetic approach involving sympathetic nerve-specific deletion of the transmembrane protein deleted in colorectal cancer (DCC). This transmembrane dependence receptor is required for postnatal development of sympathetic nerves^29^. Cell type-specific deletion of the gene encoding *Dcc* in sympathetic neurons is known to induce a severe and selective reduction in sympathetic innervation during arterial development^14,29^ but has not been evaluated in situations of visceral organ repair in adult animals.

To investigate direct interactions between sympathetic nerves and their target cells in pulmonary fibrosis, we engineered mice with *Dcc* deletion confined to TH+ sympathetic nerves (*Th-Cre; Dcc^f/f^*, Fig. 2K). In response to bleomycin, BAL noradrenaline was significantly suppressed in *Th-Cre; Dcc^f/f^* mice, compared to wild-type mice (Fig. 2L). Notably, the lack of sympathetic innervation did not affect the white blood cell count in BAL (Fig. 2M). However, it resulted in an approximately 45% reduction in collagen levels (Fig. 2N) and improved the appearance of alveolar regions in trichrome-stained tissues (Fig. 2O-R). The results provide compelling evidence that sympathetic innervation plays a direct and functional role in experimentally induced lung fibrosis.

### Loss of function of noradrenaline receptors, but not transporter, mitigates bleomycin induced lung fibrosis

Sympathetic nerve-derived noradrenaline may influence fibrosis through distinct mechanisms including neurotransmitter functions mediated by postsynaptic G-protein coupled receptors (GPCRs), and/or cellular perturbations driven by noradrenaline transporters such as solute carrier family 6 member 2 (Slc6a2^30,31^). To distinguish between these possibilities, loss-of-function studies were conducted to probe noradrenaline’s well-characterized receptors and transporters using robust pharmacological inhibitors and genetic deletion. In studies targeting GPCRs, bleomycin-challenged mice received systemic administration of the long acting, nonselective α1-adrenoreceptor (α1-AR) antagonist terazosin (Fig. 3A-D), the β1-adrenoreceptor antagonist atenolol, and the β2-adrenoreceptor antagonist ICI118,551 (Fig. 3E-H). All interventions were sufficient to improve collagen measurements and histology, supporting a role for noradrenaline’s neurotransmitter function via GPCRs in fibrosis. Conversely, fibrotic endpoints remained unchanged in noradrenaline transporter loss-of-function achieved through administration of nisoxetine, a specific inhibitor of Slc6a2 (Fig. 3I-L), and genetic approaches in *Slc6a2^-/-^* mice (Fig. 3M-O). These findings underscore the pivotal role of noradrenaline and its GPCR-mediated neurotransmitter functions in experimentally induced fibrosis.

**Fig. 3:**
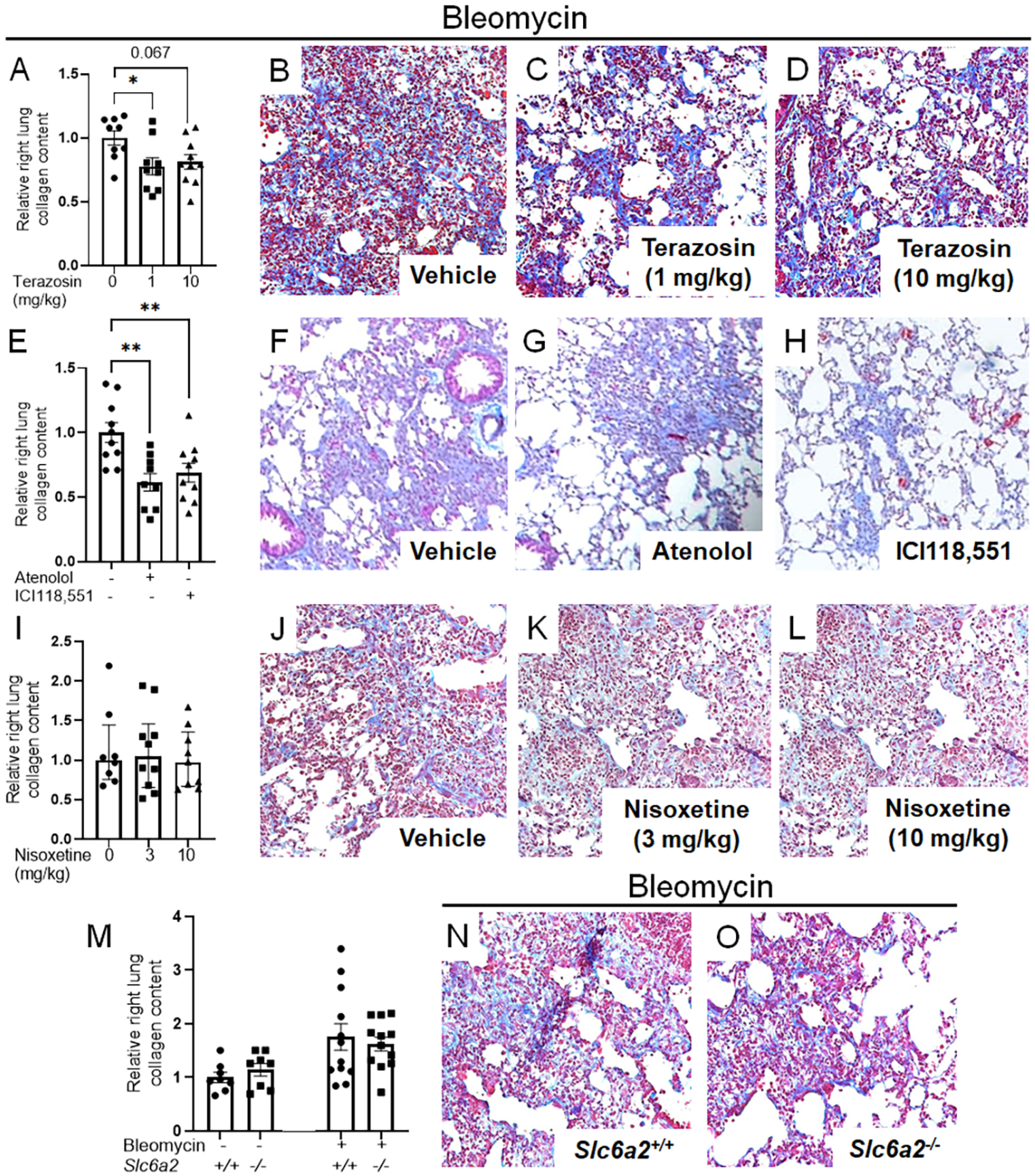
Noradrenaline-driven alveolar fibrosis requires functional neurotransmitter receptors. (A-D) Wild-type mice received orally administered terazosin, an α1 blocker, at 1 or 10 mg/kg, or vehicle from Days 5 to 13 post-bleomycin challenge and were euthanized on Day 14. Terazosin dosed at 1 mg/kg reduced collagen accumulation (A; *P* = 0.0291) and trichome staining (B-D). (E-H) Treatment with atenolol, a β1 blocker, and ICI118,551, a β2 blocker, at 1 mg/kg intraperitoneally improved both collagen deposition (E; *P* = 0.0021; *P* = 0.0098) and trichrome staining (F-H). (I-L) Intraperitoneal injections of nisoxetine, a noradrenaline transporter (NAT) antagonist, at 3 or 10 mg/kg did not reduce collagen accumulation (I) or improve trichrome staining (J-L). (M-O) Wild-type (*Slc6a2^+/+^*) and NAT-deficient (*Slc6a2^-/-^)* mice were subjected to inhaled bleomycin without any protective effect against collagen accumulation in NAT-deficient mice (M) or improvement in trichrome staining (N, O). Images were captured at 20x magnification. Data are presented as mean ± SEM or median ± IQR. Statistical analyses included Student’s t-test or ANOVA with Tukey’s test for normally distributed data, and Kruskal-Wallis tests with Dunn’s test for non-normally distributed data. \**P* < 0.05, \*\**P* < 0.01. NAT, noradrenaline transporter; Slc6a2, solute carrier family 6 member 2.

### Detection of noradrenaline and ADRA1D+ alveolar myofibroblasts in two forms of human lung fibrosis

β-adrenoreceptors are essential regulators of airway physiology and cardiac function, while under normal circumstances α1-adrenergic signaling does not possess such functions. Therefore, α1-ARs are an attractive target for the development of antifibrotic therapies in the lung and other organs. In line with this notion, the use of α1 blockers, but not β blockers, has been associated with improved clinical outcomes in conditions of inflammatory fibrosis affecting the human lung^12,14,32,33^. To integrate this human biology with the findings from our current study, the simplest explanation of our results is that nerve-derived noradrenaline mediates α-SMA+ myofibroblast effector functions through direct, cell-autonomous mechanisms. This supposition requires the identification of a noradrenaline-containing lung milieu and/or expression of one or more α1-ARs on myofibroblasts in lesional lung parenchyma. We evaluated this biology in two distinct forms of pulmonary fibrosis; namely, systemic sclerosis associated interstitial lung disease (SSc-ILD), a disease clinically characterized by sympathetic overactivation in the form of Raynaud’s phenomenon^34^, and idiopathic pulmonary fibrosis (IPF), a condition that has been linked to noradrenaline excess in several research studies^14,35^. In both settings, evidence of a noradrenaline-enriched microenvironment was found when BAL specimens from patients with either SSc-ILD or IPF exhibited increased noradrenaline concentrations relative to healthy controls (Fig. 4A). Evidence of α1-AR engagement was obtained when fibrotic alveolar regions of SSc-ILD and IPF lung tissues contained unique populations of α1-AR+ myofibroblasts. Specifically, normal donor human lungs contained α1-AR subtype D (ADRA1D) expressing cells in areas with the morphology of airways and vasculature (Fig. 4B, S2A-D), in rounded, immune-appearing cells (Fig. 4B and S2E), and in α-SMA+ smooth muscle cells adjacent to airways or vessels (Fig. S2F and G). These populations were also identified in SSc-ILD and IPF lung explants, where they are accompanied by unique populations of ADRA1D+ α-SMA+ myofibroblasts that localized to fibrotic alveolar regions (Fig. 4C-H). In contrast, only scant detection of ADRA1A and ADRA1B were observed (Fig. S3A and B). These findings were corroborated in publicly available single-cell RNA sequencing datasets generated from 6 SSc-ILD (GSE128169^36^) and 32 end-stage IPF (GSE136831^37^) patients. Consistent with the complex transcriptional regulation in which ligand-engaged α1-ARs suppress their own transcription, expression of ADRA1D itself was not observed (data not shown). However, expression of potentially relevant ADRA1D-related gene pathways such as vascular smooth muscle contraction (map04270), which culminates in myosin-actin interactions and the development of contractile force, was robustly detected in myofibroblasts, exceeding findings in fibroblasts and approaching the levels seen in *bona fide* contractile cells such as smooth muscle cells and pericytes (Fig.4I-L, S4A and B). These data suggest a potential role for noradrenaline and/or α1-ARs such as ADRA1D in the development and/or perpetuation of human lung fibrogenesis.

**Fig. 4:**
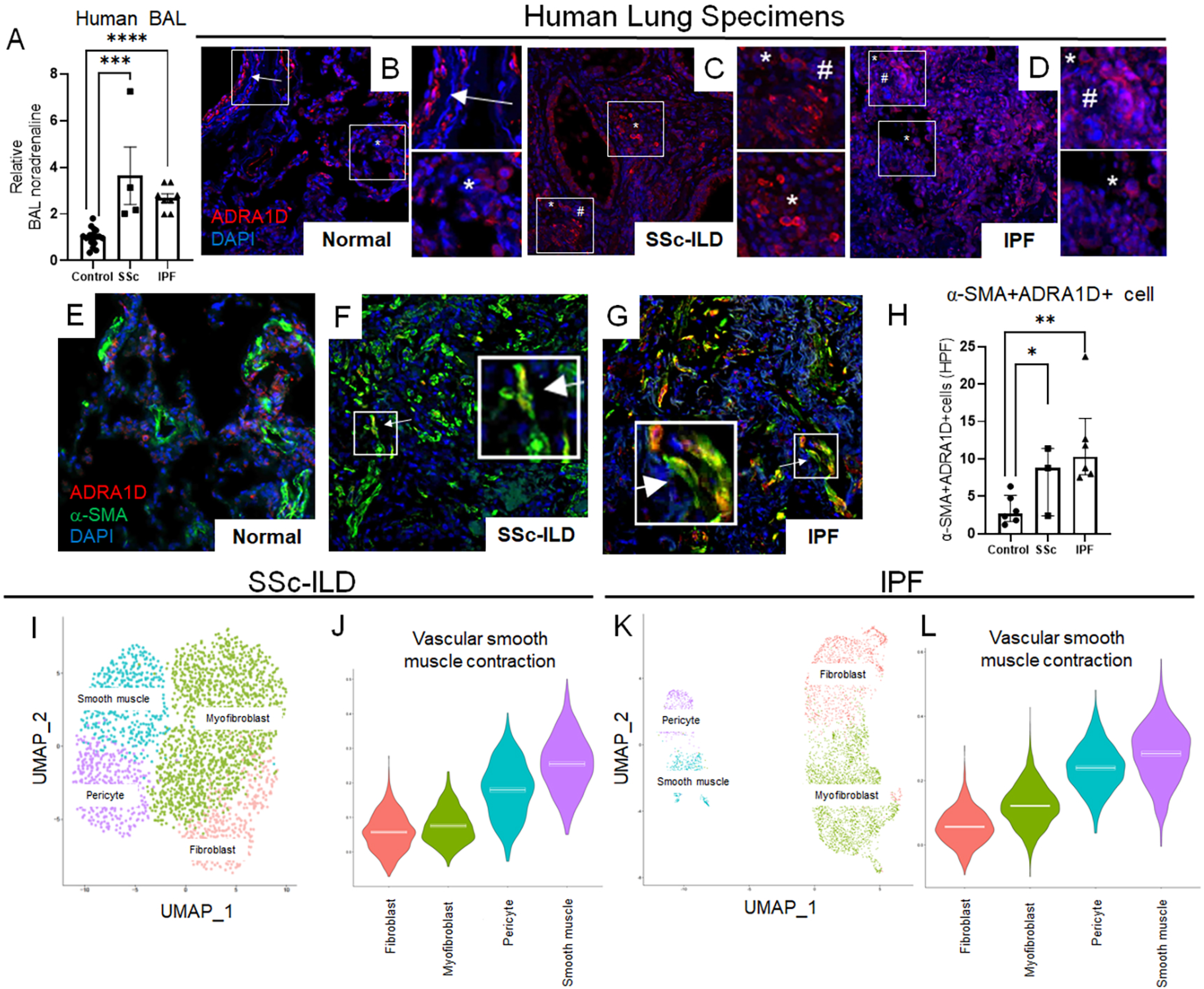
Fibrotic human lungs contain α1-adrenoreceptor-expressing myofibroblasts. (A) Relative to control, BAL noradrenaline concentrations are elevated in SSc-ILD and IPF patients (A; *P* = 0.0001 and *P* < 0.0001). (B-H) Immunofluorescence showed ADRA1D (red), α-SMA (green), and DAPI (blue) in lung tissues from healthy, SSc-ILD, and IPF patients. In normal lungs, ADRA1D expression was seen in areas with the morphology of airways and vasculature, and scattered alveolar cells (white arrows and asterisks, B). SSc-ILD and IPF lungs showed increased ADRA1D expression with additional cells in regions of fibrotic alveoli (white asterisks and hashtags, C, D). Relative to control, SSc-ILD and IPF lungs contained α-SMA+ ADRA1D+ cells in fibrotic alveoli (E-H; *P* = 0.0401 and *P* = 0.0011). (I) UMAP of single-cell RNA sequencing on stromal cell populations from the lungs of patients with SSc-ILD (n= 6) derived from GEO dataset GSE128169. (J) A violin plot demonstrated median expression of vascular smooth muscle contraction pathway across stromal cell populations in SSc-ILD lungs. (K) UMAP of single-cell RNA sequencing on stromal cell populations from the lungs of patients with IPF (n= 32) derived from GEO dataset GSE136831. (L) A violin plot demonstrated median expression of vascular smooth muscle contraction pathway across stromal cell populations in IPF lungs. Images were taken at 20x magnification; data shown as mean ± SEM or median ± IQR; analyses used Student’s t-test and Mann-Whitney test. \**P* < 0.05, \*\**P* < 0.01, \*\*\**P* < 0.001, \*\*\*\**P* < 0.0001. ADRA1D, α1-adrenoreceptor subtype D; α-SMA, alpha-smooth muscle actin; HPF, high-power field; IPF, idiopathic pulmonary fibrosis; SSc-ILD, systemic sclerosis associated interstitial lung disease; UMAP, uniform manifold approximation and projection. Red: ADRA1D. Green: α-SMA. Blue: DAPI.

### α1-adrenoreceptor antagonism induces regression of fibrotic endpoints in a next generation model of human lung fibrogenesis and in SSc-ILD and IPF lung fibroblasts

Because most patients with fibrotic conditions present with established disease, an optimally effective antifibrotic intervention should induce the suppression and/or regression of fibrotic processes. After confirming α1-AR antagonism’s benefit in a well-accepted animal model, and the existence of ADRA1D-expressing myofibroblasts in two forms of human lung fibrosis, proof-of-principle studies were conducted to determine ADRA1D’s pro-fibrotic engagement in human lungs. These studies were conducted using physiologic noradrenaline concentrations present in serum-containing conditions. Here we used a next-generation *ex vivo* modeling system in which vibratome-generated human precision-cut lung slices (PCLS) stimulated with a fibrotic cocktail permits study of fibrogenic mechanisms within the lung’s complex multicellularity and topography^38^ (Fig. 5A). In contrast to unstimulated conditions, in which detection of cells expressing α-SMA was scant (Fig. 5B), PCLS stimulated for five days with the noradrenaline containing “fibrotic cocktail” increased collagen accumulation (Fig. S5A and B), the relative expression of *ACTA2* expression in tissue lysates (Fig. S5C), and alveolar accumulation of α-SMA+ myofibroblasts (Fig. 5B-D). A significant proportion of these cells co-expressed ADRA1D (Fig. 5C and E). A fibrosis-promoting role for α1-ARs such as ADRA1D was identified when therapeutic administration of terazosin delivered after 5 days of fibrotic cocktail improved trichrome staining, reversed the alveolar accumulation of α-SMA+ fibroblasts (Fig. 5F-H), and reduced the relative expression of *ACTA2* in tissue lysates (Fig. 5I). The therapeutic potential of α1-AR antagonism was further supported when terazosin treatment suppressed the spontaneous expression of *ACTA2* in SSc-ILD lung fibroblasts (Fig. 5J). Similar but not identical results were observed in IPF, with the majority of cell lines tested showing a strong and significant reduction in the relative expression of *ACTA2* (Fig. 5K, S6). These data frame α1-AR antagonism as an antifibrotic intervention that can induce myofibroblast regression in fibrotic human lungs.

**Fig. 5:**
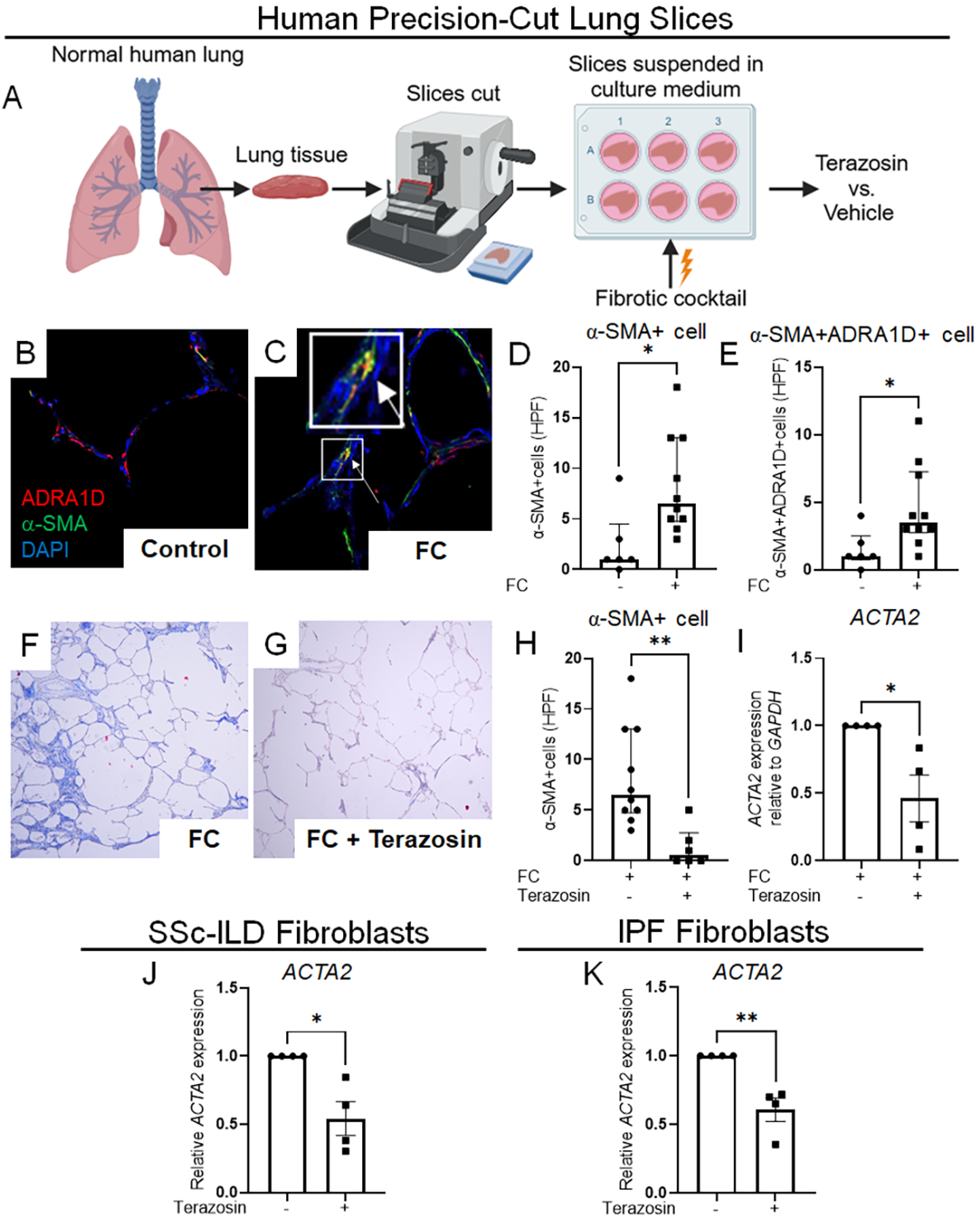
α1-adrenoreceptor antagonism induces regression of fibrotic endpoints in a next generation model of human lung fibrogenesis and in SSc-ILD and IPF lung fibroblasts. (A-I) Normal human precision-cut lung slices (PCLS) were suspended in culture media where these slices were stimulated by a fibrotic cocktail (FC), followed by treatment with terazosin or vehicle control (A). FC increased the number of cells expressing α-SMA, some of which also co-expressed ADRA1D (white arrow, B-E; *P* = 0.0106, *P* = 0.0232). Terazosin treatment improved FC-induced trichrome staining (F, G), and decreased α-SMA cells (H; *P* = 0.0017) and *ACTA2* expression in tissue lysates (I; *P* = 0.0213). (J, K) Terazosin treatment decreased *ACTA2* expression in SSc-ILD lung fibroblasts (J; *P* = 0.0345) and IPF lung fibroblasts (K; *P* = 0.0036). Images were taken at 20x magnification; data shown as mean ± SEM or median ± IQR; analyses used Student’s t-test or Mann-Whitney test. \**P* < 0.05, \*\**P* < 0.01. ADRA1D, α1-adrenoreceptor subtype D; FC, fibrotic cocktail; HPF, high-power field; IPF, idiopathic pulmonary fibrosis; PCLS, precision-cut lung slice; SSc-ILD, systemic sclerosis associated interstitial lung disease.

### Therapeutic deletion of Adra1d in myofibroblasts mitigates experimentally induced lung fibrosis

The preponderance of data suggests that interactions between sympathetic nerve-derived noradrenaline and ADRA1D+ expressing myofibroblasts sustain established fibrotic pathology in adult lungs. Given the central role of noradrenaline and adrenoreceptors in organismal homeostasis, we reasoned that the fibrogenic interaction between sympathetic nerve derived-noradrenaline and ADRA1D+ myofibroblasts must be functionally restricted to cellular perturbations encountered in the fibrotic milieu. In this paradigm, cells that express ADRA1D in uninjured conditions do not contribute to disease. To explore this idea, we characterized the kinetics of ADRA1D expression by myofibroblasts (identified by α-SMA) during bleomycin challenge and found time-dependent induction that peaked at day 14 (Fig. 6A). Based on this observation, we surmised that ADRA1D expression by the α-SMA+ myofibroblasts that accumulate during alveolar fibrosis facilitates the reception of fibrogenic noradrenergic inputs in the lung.

**Fig. 6:**
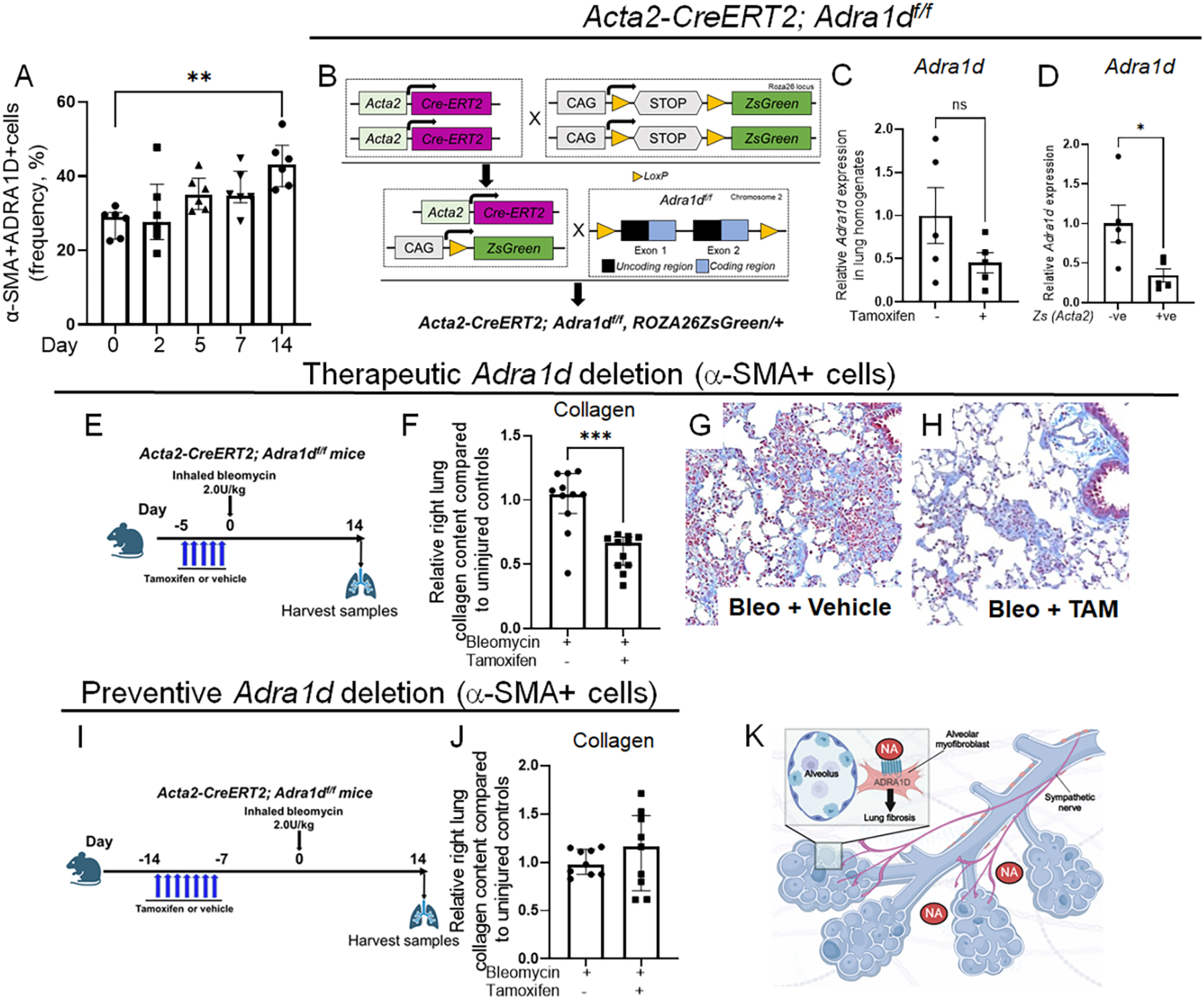
Therapeutic deletion of *Adra1d* in myofibroblasts attenuates alveolar fibrosis. (A) Following bleomycin, a time dependent increase in ADRA1D expression was observed in cells expressing α-SMA (A; *P* = 0.0026). (B) Genetic construct of *Acta2-CreERT2; Adra1d^f/f^* mice crossed with the *ROSA26^ZsGreen/+^* reporter strain. (C, D) Assessment of *Adra1d* expression in lung homogenates (C) and sorted *Zs-* and *Zs+* cells showed that *Adra1d* is specifically reduced in *Zs+* cells (D; *P* = 0.0150). (E-H) When tamoxifen was administered to delete *Adra1d* in *Acta2+* cells during active fibrogenesis (therapeutic schedule), collagen deposition was decreased (*P* = 0.0024) and trichrome staining showed reduced alveolar fibrosis. (I, J) When tamoxifen was administered to delete *Adra1d* in *Acta2+* cells before bleomycin exposure (preventive), collagen deposition was unaltered. (K) A functional interaction between sympathetic nerve-derived noradrenaline and ADRA1D+ myofibroblasts contributes to alveolar fibrosis. Images were captured at 20x magnification. Data are presented as mean ± SEM or median ± IQR with statistical analyses using the Student’s t-test, the Mann-Whitney, or Kruskal-Wallis tests. \**P* < 0.01, \*\**P* < 0.01, \*\*\**P* < 0.001. α-SMA, alpha-smooth muscle actin; NA, noradrenaline; TAM, tamoxifen; Zs, ZsGreen.

To test this hypothesis, we disrupted ADRA1D receptor function in myofibroblasts by creating a model of pan-myofibroblast *Adra1d* deletion. This objective was achieved by crossing mice with the tamoxifen-inducible *Acta2-CreERT2* promoter with a newly invented transgenic model in which exons 1 and 2 of the *Adra1d* gene were floxed (Fig. 6B, *Adra1d^f/f^*). The *Acta2-CreERT2* mouse line has been extensively utilized in studies of fibrosis^39^, vascular biology^40^, and wound healing^41^. *Acta2-CreERT2; Adra1d^f/f^* mice were viable, fertile, and displayed no significant changes in lung appearance and compared to their *Acta2-CreERT2; Adra1d^+/+^* littermates (data not shown). Tamoxifen-induced recombination efficiency was excellent, as studies performed in *Acta2-CreERT2; Adra1d^f/f^* crossed with the *ROSA26^ZsGreen/+^* reporter strain showed that *Adra1d* expression was specifically reduced in cells with an active *Acta2* promoter (Fig. 6C and D, S7).

Then, to illustrate the concept that ADRA1D allows α-SMA+ effector cells to receive fibrosis-promoting noradrenergic signals, we delivered tamoxifen according to a schedule timed to delete *Adra1d* in the myofibroblasts that emerge during alveolar fibrogenesis (Figure 6E). In this system, relative to corn oil, the lungs of tamoxifen-treated *Acta2-CreERT2; Adra1d^f/f^* mice exhibited improvements in total lung collagen content and alveolar fibrosis shown by trichrome staining (Fig. 6F-H, S8A). Conversely, tamoxifen administered before injury (which targets *Adra1d* in α-SMA+ cells present in nonfibrotic conditions) showed no such benefit (Fig. 6I and J, S8B). These findings provide *in vivo* evidence of a functional interaction between nerve-derived noradrenaline and ADRA1D that is pivotal for this fibrogenic axis comprised of sympathetic nerves and myofibroblast effector cells (Fig.6K).

## DISCUSSION

The discovery that sympathetic nerves orchestrate pulmonary fibrosis by directing myofibroblast function provides fundamental new insights into the repair and pathologic remodeling of injured organs. Specifically, we provide compelling evidence of a mechanism though which sympathetic innervation controls noradrenergic fibrotic remodeling in settings of maladaptive repair in the lungs of adult mammals. This functional unit is conserved across species (mouse and human) and diseases (SSc-ILD and IPF). Our work is a major advance over prior studies due to its state-of-the-art methods including light sheet imaging, nerve-fibroblast co-culture, cell specific knockout mice, well accepted pharmacologic inhibitors, next generation *ex vivo* fibrosis models, and specimens from two distinct forms of human lung fibrosis. These technical achievements facilitated novel observations of aberrantly patterned TH+ nerves in distal airways, functional benefit enacted by direct loss of sympathetic nerves and pharmacologic interruption of noradrenaline’s neurotransmitter receptors, and the evidence of functional noradrenergic input from sympathetic nerves to ADRA1D+ myofibroblasts in well accepted mouse models, advanced lung mimetics, and several fibrotic conditions affecting human lungs. Because the lung depends upon autonomic innervation for proper development and functional homeostasis^22^, we predict that this newly described nerve-fibroblast axis controls numerous aspects of lung physiology in health and disease. Furthermore, considering the pivotal and highly conserved nature of sympathetic innervation, noradrenaline, and myofibroblasts across tissues, we predict that this interaction underpins homeostasis, repair, and pathologic remodeling in numerous organs.

The discovery of a functional axis involving sympathetic nerves and fibrogenic myofibroblasts augments the nascent field of nerve-lung interactions in the adult mammals. In recent years the availability of advanced imaging methods^26^, genetic tools^42^, and single-cell sequencing^37^, has facilitated studies of autonomic and/or sensory innervation as contributors to lung pathology. Much of this work provided indirect evidence, and none studied myofibroblast innervation in alveolar repair and fibrosis. Therefore, our use of genetic models and neuronal cell culture to show that sympathetic nerves are absolutely required for maximal fibrosis provides first of its kind insights to the neurobiology of alveolar repair and remodeling that raise numerous questions that warrant further study. For example, the nature of nerve endings has yet to be determined, as it is not clear whether sympathetic nerve-derived neurotransmitters such as noradrenaline reach their target through free nerve endings, diffusion, or alternate mechanisms^43^. Additionally, the roles of non-adrenergic nerve functions and compensatory mechanisms, such as denervation hypersensitivity^44^, remain elusive. Further, since *Th-Cre* drivers and pharmacological interventions are not lung-specific, we cannot rule out the possibility of contributions from mechanisms occurring in other peripheral organs or the central nervous system. Rather than undermining our findings, this latter possibility bolsters the exciting concept of systemic control of tissue level responses and provides an urgent mandate for additional study of this intriguing new paradigm.

Our study provides definitive evidence of a new fibrotic pathway involving sympathetic nerves and myofibroblasts, the mechanism of which is currently unknown. The benefit was observed only when *Adra1d* deletion occurred in myofibroblasts during fibrogenesis, suggesting that the mechanism does not involve the expansion of an α-SMA+ ADRA1D+ progenitor population present within smooth muscle cells or alveolar myofibroblasts. Instead, it appears to be an effect on fibrogenesis mediated by an emerging population of ADRA1D+ myofibroblasts. As shown by our *ex vivo* studies, this process is likely at least partially due to sympathetic innervation-induced alterations in the decisions that myofibroblasts make regarding their functional state. Additionally, although our findings provide clear evidence that myofibroblasts are a primary recipient of cues from sympathetic nerves, we cannot exclude the possibility of non-cell-autonomous mechanisms. In fact, the widespread expression of receptors for sympathetic nerve-derived neurotransmitters makes such a possibility highly likely and provides numerous new avenues for investigative studies aimed at mechanistic discovery. Finally, the existence of numerous pharmacologic agents targeting noradrenergic GPCRs and the emergence of clinical strategies based on targeted neuromodulation^45^ leave us well positioned for therapeutic breakthroughs.

In conclusion, we provide evidence of a functional axis involving sympathetic nerves and myofibroblasts in mammalian lung fibrosis. This discovery provides new information about how cells send and receive signals in the adult lung. It also provides new insight into how nerves communicate with tissues during conditions of injury and regenerative failure. It shows how two distinct organs – autonomic nerves and the lung – interact during fibrogenesis in adult mammals. Last, it provides proof-of-concept evidence for the development of neuromodulation-based strategies to treat myofibroblast driven conditions in humans. Further study of these important areas will illuminate paradigm shifting discoveries in tissue repair with the goal of preventing and perhaps reversing fibrosis and advancing human health.

## MATERIALS AND METHODS

### Study approval

All animal experiments were approval by the Yale University Institutional Animal Care and Use Committee (IACUC) and were conducted in compliance with the guidelines outlined in the Guide for the Care and Use of Laboratory Animals^46^. Banked, archived bronchoalveolar lavage fluid specimens (BALs) from control subjects, systemic sclerosis associated interstitial lung disease (SSc-ILD) and IPF subjects were provided by the Yale Lung Repository and the Yale Scleroderma Repository under exemption 8. De-identified lung tissues had been previously obtained at the time of lung transplantation under a human subjects protocol approved by the University of Pittsburgh IRB. Additional samples were sourced from de-identified clinical specimens provided by the Yale Pathology Biobank.

### Animals

Wild-type male and female C57BL/6J mice obtained from The Jackson Laboratory (Bar Harbor, ME) were used between 8 to 12 weeks of age. Transgenic mice used in this study included *B6.Cg-7630403G23Rik^Tg(Th-cre)1Tmd^/J* (*Th-Cre*), *B6(Cg)-Tg(Acta2-cre/ERT2)1Ikal/J* (*Acta2-CreERT2*), *B6.129P2(Cg)-Slc6a2^tm^*^1^*^.1Hhg^/J* (*Slc6a2^-/-^)*, and *B6.Cg-Gt(ROSA)26Sor^tm^*^6^*^(CAG-ZsGreen^*^1^*^)Hze^/J* (*ROSA26^ZsGreen/+^*) mice and were all purchased from The Jackson Laboratory (Bar Harbor, ME). *C57BL/6JGpt-Dcc^em1Cflox^/Gpt* (*Dcc^f/f^*) mice and *B6/JGpt-Adra1d^em1Cflox^/Gpt* (*Adra1d^f/f^*) mice were purchased from GemPharmatech Co., Ltd, Nanjing, China. All mice were on the C57BL/6J background.

### Inhaled bleomycin administration

Oropharyngeal bleomycin administration (2.0U/kg, Mckesson, 63323013610) was performed as previously reported^12,14^. Control and treatment groups were maintained under identical conditions. Specimens from animals that did not survive to the prespecified endpoints were excluded from analysis.

### Administration of experimental agents

Mice were allocated into groups that received daily intraperitoneal injections of PBS control or experimental agents from days 5-13 post bleomycin. Experimental agents included atenolol (1 mg/kg, Sigma-Aldrich, A7655), ICI118,551 (1 mg/kg, Sigma-Aldrich, I127), or nisoxetine (3 or 10 mg/kg, MilliporeSigma, 57754-86). An additional cohort received PBS or terazosin (1 or 10 mg/kg, U.S. Pharmacopeia, 1643452) via daily oral gavage on the same schedule.

### Tamoxifen Administration

Depending on the deletion schedule, mice received injections of tamoxifen (2 mg per day for either 5 or 7 days, Sigma-Aldrich, T5648) or an equivalent volume of corn oil as a vehicle control. Thereafter, mice either received immediate bleomycin inhalations or were allowed a 7-day rest period before bleomycin administration.

### Sacrifice and lung harvest

Fourteen days following bleomycin, mice underwent terminal anesthesia, bronchoalveolar lavage (BAL), median sternotomy, right heart perfusion with 1x PBS, and *en bloc* lung resection^12,47^.

### Bronchoalveolar lavage cell quantification

BAL cell counts were determined as previously described^48^.

### Determination of noradrenaline concentrations

Noradrenaline concentrations were quantified in plasma, BAL fluid, and cell culture supernatants using high-sensitivity ELISA kits (DLD Diagnostika, GMBH, NOU39-K01) as previously reported^12,14^.

### Collagen quantification

Lungs were snap frozen in liquid nitrogen and stored at −80°C until quantification of lung collagen using the Sircol Collagen Assay (Biocolor Ltd., S1000) or Hydroxyproline Assay (QuickZyme Biosciences, QZBHYPRO5) as previously reported^12,14,47^.

### Flow cytometry analysis on digested lung tissues

Flow cytometry analysis was carried out using an LSRII flow cytometer (BD Biosciences, Franklin Lakes, NJ) on lung tissue suspensions from euthanized mice. The lungs were first perfused with 1x PBS, then harvested, minced, and digested in 1x PBS containing 150 μg/mL collagenase (MilliporeSigma, C5138) and 20 U/mL DNase I (Roche, 04716728) for 1 hour at room temperature. The digestion was quenched with 1x PBS. Suspensions were filtered through a 40-μm cell strainer and centrifuged at 495 g for 10 minutes at 4°C. The supernatants were discarded, and the pellets were resuspended in 10 mL of 1x PBS for cell counting. After another round of centrifugation, cell pellets were resuspended in FACS buffer (1× PBS with 1% fetal bovine serum [FBS], 0.01% NaN3, and 1 mM EDTA) to a concentration of 1 x 10^6^ cells/mL. For staining, 1 x 10^6^ cells were incubated with a series of antibodies at specific dilutions in FACS buffer containing 10% normal goat serum (NGS) for 1 hour at 4°C. The staining panel included: rat anti-CD45 FITC (1:100, eBioscience, 11-0451-82), rat anti-CD45 PE (1:100, BD Biosciences, 553081), rat anti-CD45 PerCP (1:1000, eBioscience, 45-0451-82), rat anti-CD45 APC (1:100, eBioscience, 17-0451-82) for compensation controls, rabbit anti-ADRA1D (1:100, Abcam, ab84402), and mouse anti–α-SMA (1:250, intracellular staining, Abcam, ab7817). Secondary antibodies were used as necessary for detection of unconjugated primary antibodies. Following staining, cells were washed, filtered through a 40-μm cell strainer, and then subjected to data acquisition on the LSRII flow cytometer using FACSDiva software (BD Biosciences, Franklin Lakes, NJ). Data analysis was conducted using FlowJo software (BD Biosciences, Franklin Lakes, NJ), employing gating strategies refined by control samples incubated without primary antibodies. This approach ensured accurate identification of positive and negative cell populations. *ZsGreen*-expressing and non-expressing cells were isolated from *Acta2-CreERT2; Adra1d^f/f^, ROSA26^ZsGreen/+^* mice, that received tamoxifen and bleomycin, through fluorescence-activated cell sorting (FACS).

### Histologic analysis

Resected left lungs were fixed in 10% formalin, embedded in paraffin, sectioned, and stained with Masson’s Trichrome^12^. Precision-cut lung slices were also stained with Masson’s Trichrome.

### Sympathetic nerve harvesting and culture

Neuronal cells were harvested from superior cervical ganglion of neonatal mice on postnatal days 1-3 that were incubated in digestion buffer, which comprised collagenase type II (Millipore Sigma, C2-28-100mg) and dispase type II (Millipore Sigma, D4693-1G), for 60 minutes at 37°C. Following incubation, careful mechanical dissociation was performed using a 10 µL long-tip pipette in a dissection solution comprised of Leibovitz’s L-15 medium and fat-acid-free BSA (400 mg/500 mL). After centrifugation at 2,000 RPM for 5 minutes, the cells were resuspended in glutamine-supplemented Dulbecco’s Modified Eagle Medium (DMEM)/F-12 containing 10 ng/mL nerve growth factor (NGF, Thermo Fisher, 13257-019) and plated on poly-D-lysine hydrobromide (MP Biomedicals LLC, 0210269480) -precoated 24-well plates (seeding nerve cells from two ganglia per well).

### Sympathetic nerve + normal human lung fibroblast (NHLF) co-culture

After a growth period of 7 days, sympathetic neuronal cells were co-cultured with NHLFs for 48 hours and incubated with the profibrotic stimulus TGFβ1 (0 or 5 ng/mL). Normal human lung fibroblasts were procured from Lonza (Allendale, NJ). Cells used at passages 5-10 were cultured to confluence in Dulbecco’s Modified Eagle Medium/10% Fetal Bovine Serum/1% penicillin-streptomycin. Approximately 20,000 cells were seeded into each well of 24-well plates. After co-culture, NHLFs were collected for real time quantitative PCR analysis.

### Precision-cut lung slices

Normal human precision-cut lung slices (PCLS) were purchased from the Institute for In Vitro Sciences (Gaithersburg, MD). PCLS were rapidly thawed and transferred to an acclimation medium, which consisted of DMEM/F12 (Thermo Fisher, 11320033), 0.2% Primocen® (Invivogen, ANT-PM-1), 1% insulin-transferrin-selenium (Thermo Fisher, 41400045), 1% Antibiotic Antimycotic solution (Millipore Sigma, A5955), 2 µM hydrocortisone (Millipore Sigma, H0888), and 2-phospho-L-ascorbic acid trisodium salt (Millipore Sigma, 49752). Following three days of culture in this medium, PCLS were transferred to a culture medium composed of DMEM supplemented with 0.2% Primocen® and 1% insulin-transferrin-selenium for three days. Next, PCLS were cultured in the absence or presence of a serum-containing pro-fibrotic media for five days consisting of the following: 5 ng/mL TGFβ1 (Bio-Techne Corporation, 240-GMP-010), 5 μM platelet-derived growth factor-AB (PDGF-AB, ThermoFisher, 100-00AB-10UG), 10 ng/mL tumor necrosis factor alpha (TNF-α, R&D Systems, 210-TA), and 5 μM lysophosphatidic acid (Cayman Chemical, 62215). PCLS were then treated in the absence or presence of 10 µM terazosin (U.S. Pharmacopeia, 1643452) for 24 hours. Subsequently, the PCLS were harvested and prepared either for RNA analysis using Qiazol or for histological examination.

### Human lung fibroblast culture

SSc-ILD and IPF fibroblasts were procured from Asterand Bioscience (Detroit, MI) or provided by Dr. Carol Feghali-Bostwick (Medical University of South Carolina, SC). Cells used at passages 5-10 were cultured to confluence in Dulbecco’s Modified Eagle Medium/10% Fetal Bovine Serum/1% penicillin-streptomycin. Approximately 400,000 cells were seeded into each well of a 6-well plate. After 24 hours of serum deprivation, the cells were treated with/without terazosin (10 μM; U.S. Pharmacopeia, 1643452) for 48 hours. After treatment, samples were collected for real time quantitative PCR analysis.

### Adipo-Clear tissue clearing and three-dimensional whole mouse lung imaging

Standard intracardiac perfusion was performed using 20 mL of 1x PBS containing 10 µg/mL heparin (Sigma-Aldrich, H3393-50KU) at 4 °C until the blood was completely removed from the lung. The perfusate was then switched to 20 mL of fixative solution (4% paraformaldehyde [PFA] in 1x PBS) at 4 °C, and the lung was post-fixed in 4% PFA in 1x PBS overnight at 4 °C. After washing in 1x PBS/0.02% NaN3 (shaken at room temperature (RT), 1h x 3 times), the sample was dehydrated in a graded series of 20%, 40%, 60%, 80%, and 100% (three times) methanol in 0.05x PBS buffer for 60 minutes at each step. The sample was delipidated with 100% dichloromethane (DCM, Sigma-Aldrich, 270997-12X100ML) (three times) with incubation times of 2 hours, overnight, and 1 hour. Then, the sample was washed three times with 100% methanol for 60 minutes each and bleached with 5% H_2_O_2_ in methanol overnight at 4 °C. The sample was rehydrated in a reversed series of methanol/0.05x PBS buffer: 80%, 60%, 40%, 20%. The sample was then washed with 5% DMSO/0.3M Glycine/PTxwH buffer (PBS/0.1% Triton X-100/0.05% Tween 20/2 µg/mL heparin) twice for 1 hour and 2 hours each at RT, followed by washes in PTxwH: shaken at RT, 60min x 3. The sample was incubated with primary antibodies (sheep anti-tyrosine hydroxylase, 1:500, Millipore Sigma, AB1542, and mouse anti-α-SMA, 1:500, Abcam, ab7817) diluted in PTxwH (with 1% donkey serum) and shaken at 37°C for 10 days. The sample was then washed in PTxwH, shaken at 37°C for 1 hour twice and 2 hours twice, followed by an overnight wash. Subsequently, the sample was incubated with secondary antibodies diluted in PTxwH (with 1% donkey serum) and shaken at 37°C for 7 days. The sample was washed with PTxwH buffer through a series of incubation steps: 1 hour twice, 2 hours twice, overnight, and for 1 day. This was followed by a wash with 1x PBS, also in a series of incubation steps: 1 hour twice, 2 hours twice, and overnight. Then, the sample was dehydrated in a methanol gradient with 20%, 40%, 60%, 80%, and 100% methanol (three times) for 60 minutes each. The sample was then incubated in 100% DCM for 1 hour with shaking (twice). Finally, the sample was incubated in dibenzyl ether (DBE, Sigma-Aldrich, 108014-1KG) overnight with mild shaking to achieve refractive index matching and kept in DBE until 3-dimensional mouse lung imaging was performed with UltraMicroscope Blaze (Miltenyi Biotec, Bergisch Gladbach, Germany). Imaris 10.0 software (Oxford Instruments, Abingdon, United Kingdom) was used for the analyses.

### Immunofluorescence methods

Immunofluorescence analysis for the detection of α1-adrenoreceptors (α1-AR) and α-SMA in human lung sections and mouse sympathetic nerve cells utilized the following primary antibodies: rabbit anti-ADRA1A (1:100, Abcam, ab137123), rabbit anti-ADRA1B (1:250, Abcam, ab169523), rabbit anti-ADRA1D (1:250, Abcam, ab84402), rabbit anti-ADRA1D (1:250, LSBio, LS-A12-50), mouse anti–α-SMA (1:250, Abcam, ab7817), and rabbit anti-tyrosine hydroxylase (1:250, Abcam, ab112). After primary antibody application, sections were subjected to secondary antibody detection and nuclei were counterstained with 4′,6-diamidino-2-phenylindole (DAPI). Human prostate tissues were employed as positive controls for the specificity of the anti-ADRA1D antibody. Negative controls included slides processed without primary antibodies. Imaging was performed using a Nikon Eclipse microscope (Nikon Corporation, Tokyo, Japan), equipped with coherent 488 and 561 nm lasers. Image capture was facilitated by an Andor iXON3 EMCCD detector, with NIS Elements AR software (Nikon Corporation, Tokyo, Japan) controlling the imaging process.

### RNA isolation and real time quantitative PCR

Total cellular RNA was extracted using the miRNeasy Mini-Kit (Qiagen, 217084) according to the manufacturer’s protocol. This RNA was then reverse-transcribed using the Power SYBR™ Green RNA-to-CT™ 1-Step Kit (Applied Biosystems, 4389986). Subsequent analysis focused on the expression levels of *ACTA2* and *GAPDH* (human) and *Adra1d* and *Actb* (mouse) using specific primers for each gene on the ViiA 7 Real-Time PCR System (Thermo Fisher, Waltham, MA). Relative gene expression was quantified using the 2-delta Ct method, consistent with our methodologies^47^. *ACTA2* expression from all PCR replicates was reported in experiments co-culturing nerve cells and NHLFs.

Human primers were:

*ACTA2*-F: 5′ -GTGTTGCCCCTGAAGAGCAT -3′;
*ACTA2*-R: 5′ -GCTGGGACATTGAAAGTCTCA -3′;
*GAPDH*-F: 5′ -TGGAGAAGGCTGGGGCTCATTT-3′;
*GAPDH*-R: 5′ -TGGTGCAGGAGGCATTGCTGAT-3′;

Mouse primers were:

*Actb*-F: 5′ -GGCTGTATTCCCCTCCATCG-3′;
*Actb*-R: 5′ -CCAGTTGGTAACAATGCCATGT-3′;
*Adra1d*-F: 5′ -AATCTTGCTGCACTAGGGCTCT-3′;
*Adra1d*-R: 5′-CTAGTCATGTCAACAGGAGCTGGA-3′;

### Single-cell RNA sequencing (scRNA seq) analysis

Analyses of publicly available scRNA seq datasets (GSE128169 for SSc-ILD and GSE136831 for IPF) were performed with Seurat version 5.0.318. Seurat object were created for each dataset and data was normalized. Mitochondrial and cell cycle genes were identified and used for adjustments. RNA seq annotation was performed with Azimuth^49^. Principal components analysis was performed, and 70 significant principal components were chosen using Jack Straw for downstream cell clustering and visualization using the uniform manifold approximation and projection (UMAP) technique. To identify cell types expressing genes associated with ADRA1D-related pathways, the AddModuleScore, a Seurat function that calculates the gene expression level of a cluster of genes on a single-cell level and then subtracts the aggregated expression of a randomly assigned control set, was used.

### Graphics

Graphics were designed using BioRender.com (Toronto, Ontario, Canada).

### Statistical Analyses

All data are presented as mean ± SEM or median ± IQR unless stated otherwise. Normally distributed data were compared using 1- or 2-tailed student’s t test or ANOVA with Tukey’s multiple comparisons test. Non-normally distributed data were compared using the nonparametric Mann-Whitney test or Kruskal-Wallis test with Dunn’s multiple comparisons test. GraphPad Prism 9.0 (GraphPad Software, CA) was used for all these analyses. A p-value < 0.05 corrected for multiple testing was considered significant.

## Acknowledgements

GI conceptualized the project, administered its execution, secured funding, conducted the investigation, interpreted the data, and contributed to both writing and editing the manuscript. XP, AG, SS, AJ, JM, JZ, SW, DO, SY, CJL, AL, TS, BH, YS, RG, RH, HS, and JLG carried out the investigation and data analysis. KIJ administered the project. MH and CFB provided resources. TSS and MS contributed to project conceptualization. CR provided supervision, resources, administered the project, and acquired funding alongside ELH, who contributed similarly in these areas. ELH also interpreted the data, assisted in conceptualization and participated as the corresponding author in writing and editing the manuscript. All authors were involved in preparing the manuscript and approved the final version for submission. We extend our heartfelt thanks to all the patients living with fibrotic lung disease. Your strength, courage, and resilience deeply inspire us and fuel our commitment to advancing research in this field.

## Data Availability

The data that support the findings of this study are available on request from the corresponding author.

