## Supplemental Figures for "A Nerve-Fibroblast Axis in Mammalian Lung Fibrosis"

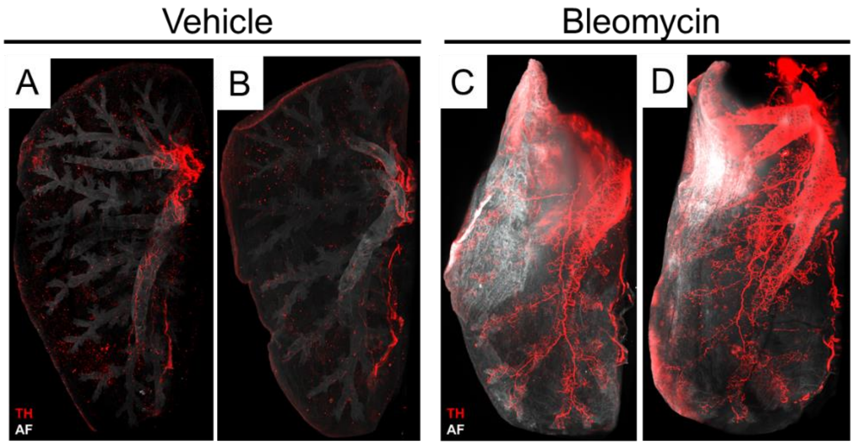


**Fig. S1: 3D visualization of sympathetic innervations in murine lungs.** (A-D) Light-sheet microscopy visualized sympathetic innervations in the entire murine lungs immunolabeled with anti-TH (red). 3D projections in uninjured lungs demonstrated that TH+ nerves were localized to proximal airways, with minimal extension into distal airways (A and B depict two replicates to complement the images in Figure 1). In fibrotic lungs, TH+ nerves extended into peripheral airways (C and D depict two replicates to complement Figure 1). AF, autofluorescence; TH, tyrosine hydroxylase.

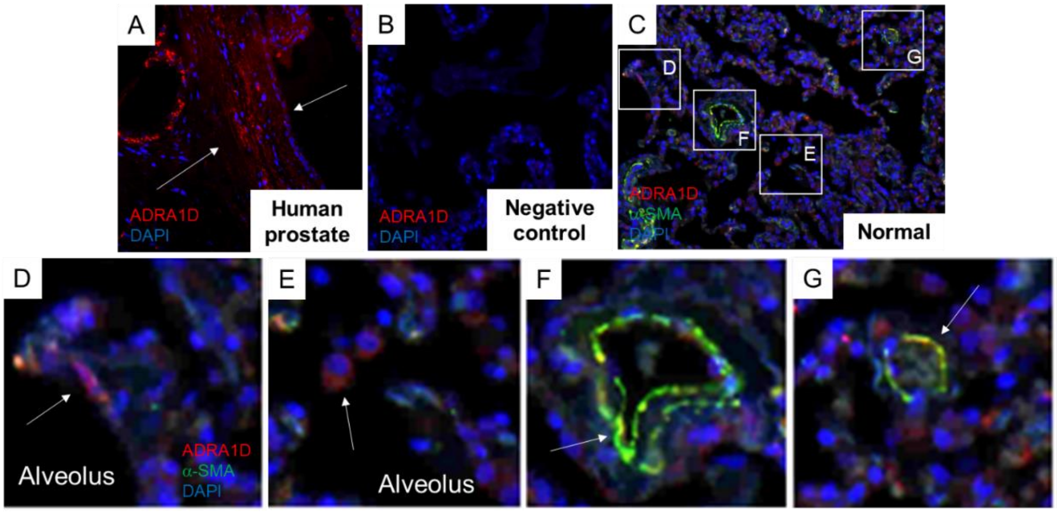


**Fig. S2: Localization of ADRA1D-expressing cells in normal human lungs.** Immunofluorescence imaging demonstrated ADRA1D (red), α-SMA (green), and DAPI (blue) staining in explant normal human lung and prostate tissues. (A) Human prostate tissue served as a positive control, showing pronounced ADRA1D signal in morphologically characteristic smooth muscle cells (white arrows). (B) Negative control without primary antibodies in normal human lung tissue. (C-G) In normal human lung tissue, ADRA1D expression was observed in cells adjacent to alveoli (white arrow, D) and in dispersed alveolar cells (white arrow, E). Additionally, ADRA1D was partially co-expressed with α-SMA-positive cells surrounding tubular structures morphologically resembling airways or vessels (white arrows, F, G). The image was captured at 20x magnification. ADRA1D, α1-adrenoreceptor subtype D; α-SMA, alpha-smooth muscle actin; DAPI, 4′,6-diamidino-2-phenylindole.

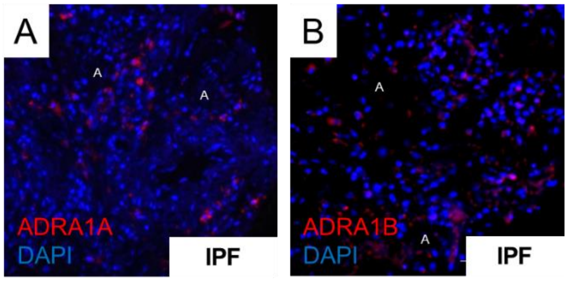


**Fig. S3: Localization of ADRA1A/ADRA1B-expressing cells in fibrotic human lungs.** (A) Minimal expression of ADRA1A near alveoli in IPF lung tissues. (B) Low expression of ADRA1B near alveoli in the same tissues. The image was captured at 20x magnification. A, alveolus; ADRA1A, α1-adrenoreceptor subtype A; ADRA1B, α1-adrenoreceptor subtype B; DAPI, 4′,6-diamidino-2-phenylindole.

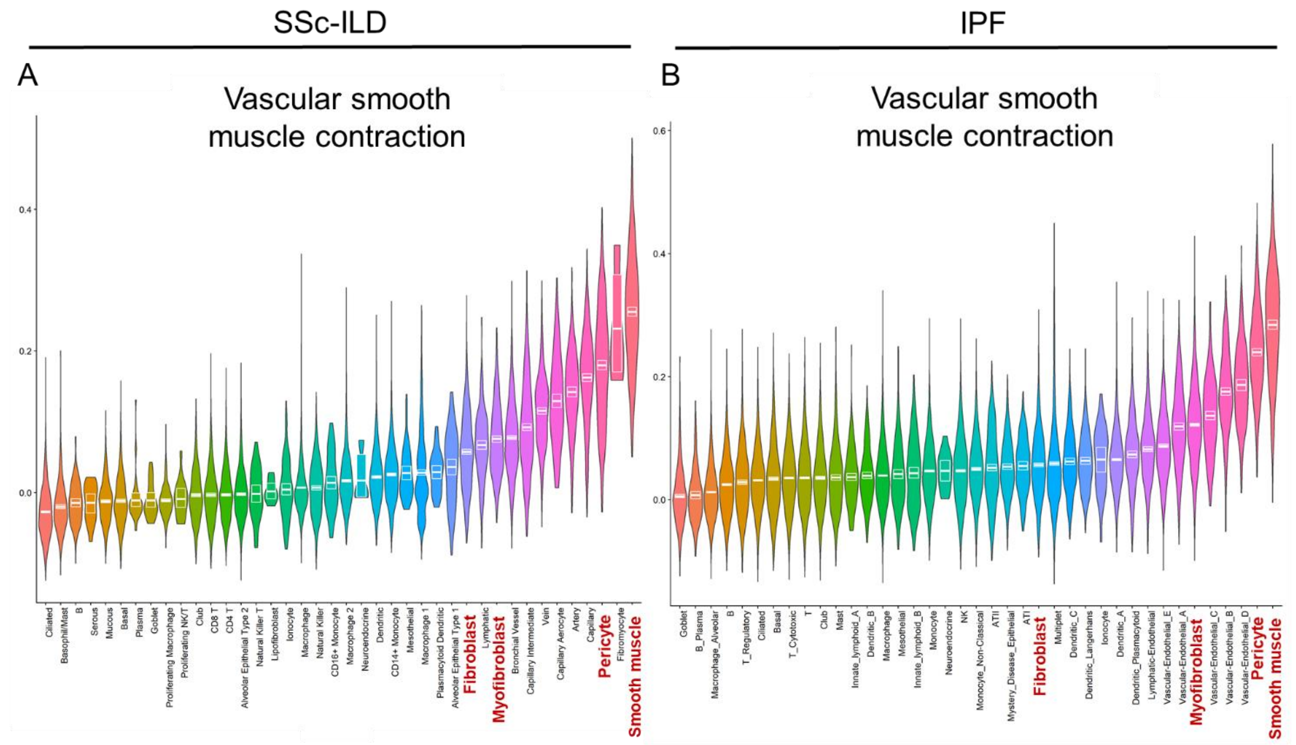


**Fig. S4: Expression of the ADRA1D-related pathway in SSc-ILD and IPF single-cell RNA Seq datasets.** Violin plots demonstrated median expression of vascular smooth muscle contraction pathway across multiple cell populations in SSc-ILD (A) and IPF lungs (B). IPF, idiopathic pulmonary fibrosis; SSc-ILD, systemic sclerosis associated interstitial lung disease. Red: stromal cells.

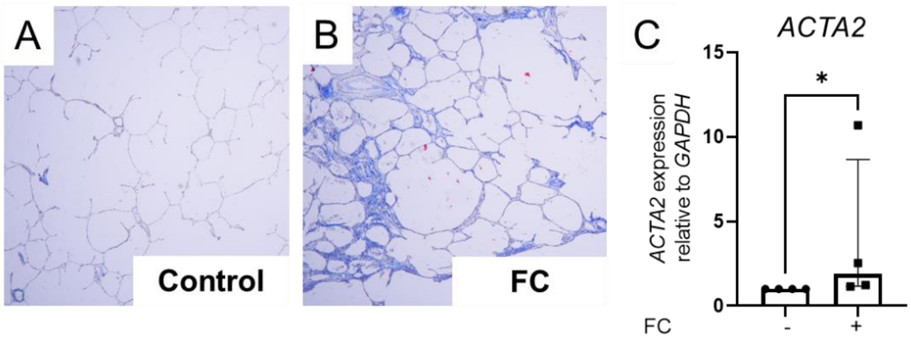


**Fig. S5: Fibrotic cocktail induces collagen accumulation and *ACTA2* expression in human PCLS.** Human PCLS stimulated with a fibrotic cocktail increased collagen content in trichrome staining (A, B) and *ACTA2* expression (C; $P = 0.0286$). Images were taken at 20x magnification; data shown as median ± IQR; analyses used Mann-Whitney test. *$P < 0.05$. FC, fibrotic cocktail; PCLS, precision-cut lung slice.

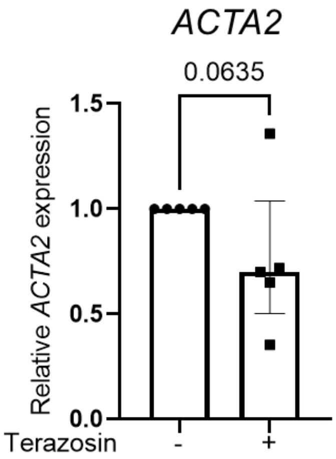


**Fig. S6: Relative expression of *ACTA2* in terazosin treated IPF lung fibroblasts.** Terazosin treatment reduced the relative expression of *ACTA2* in four out of five IPF fibroblast cell lines. Data are presented as median ± IQR. Statistical analysis used Mann-Whitney test.

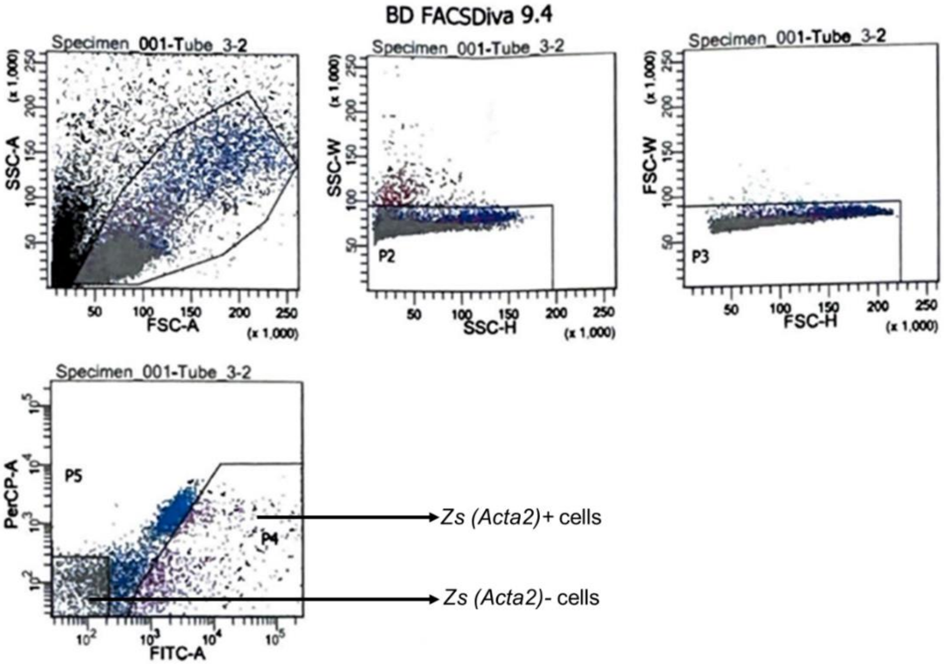


**Fig. S7: FACS gating strategy for sorting *ZsGreen* (*Acta2*)-expressing and non-expressing cells.** FACS, fluorescence-activated cell sorting.

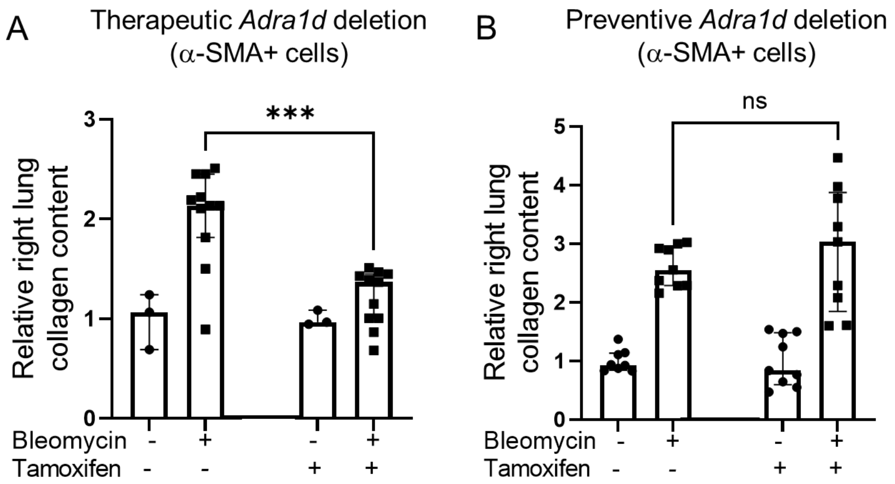


**Fig. S8: *Adra1d* deletion in α-SMA cells in bleomycin-induced lung fibrosis.** (A) In *Acta2-CreERT2; Adra1d$^{f/f}$ mice*, therapeutic tamoxifen administration to delete *Adra1d* in myofibroblasts during active fibrogenesis mitigated bleomycin induced collagen accumulation ($P$ = 0.0024). (B) When tamoxifen was administered to delete *Adra1d* in α-SMA-positive progenitor cells before bleomycin exposure (preventive), it did not reduce collagen deposition. Data are presented as median ± IQR. Statistical analyses involved the Mann-Whitney test. ***$P$ < 0.001. ADRA1D, α1-adrenoreceptor subtype D; α-SMA, alpha-smooth muscle actin.
